# Causal Effects of Blood Lipids on Amyotrophic Lateral Sclerosis: A Mendelian Randomization Study

**DOI:** 10.1101/447581

**Authors:** Ping Zeng, Xiang Zhou

**Affiliations:** Department of Epidemiology and Biostatistics, Xuzhou Medical University, Xuzhou, 221004, Jiangsu, China; Department of Biostatistics, University of Michigan, Ann Arbor, 48109, Michigan, USA; Center for Statistical Genetics, University of Michigan, Ann Arbor, Michigan, USA

## Abstract

Amyotrophic lateral sclerosis (ALS) is a late-onset fatal neurodegenerative disorder that is predicted to increase across the globe by ~70% in the following decades. Understanding the disease causal mechanism underlying ALS and identifying modifiable risks factors for ALS hold the key for the development of effective preventative and treatment strategies. Here, we investigate the causal effects of four blood lipid traits that include high density lipoprotein (HDL), low density lipoprotein (LDL), total cholesterol (TC), and triglycerides (TG) on the risk of ALS. By leveraging instrument variables from multiple large-scale genome-wide association studies in both European and East Asian populations, we carry out one of the largest and most comprehensive Mendelian randomization analyses performed to date on the causal relationship between lipids and ALS. Among the four lipids, we found that only LDL is causally associated with ALS and that higher LDL level increases the risk of ALS in both the European and East Asian populations. Specifically, the odds ratio of ALS per one standard deviation (i.e. 39.0 mg/dL) increase of LDL is estimated to be 1.14 (95% CI 1.05 - 1.24, *p* = 1.38E-3) in the European and population and 1.06 (95% CI 1.00 - 1.12, *p* = 0.044) in the East Asian population. The identified causal relationship between LDL and ALS is robust with respect to the choice of statistical methods and is validated through extensive sensitivity analyses that guard against various model assumption violations. Our study provides important evidence supporting the causal role of higher LDL on increasing the risk of ALS, paving ways for the development of preventative strategies for reducing the disease burden of ALS across multiple nations.

## Introduction

Amyotrophic lateral sclerosis (ALS) is a late-onset fatal neurodegenerative disorder that is characterized by progressive degeneration of upper and lower motor neurons in adults (1). Patients of ALS typically experience signs and symptoms of progressive muscle atrophy and weakness, dysphagia, dysarthria, and eventually respiratory failure. About 10% of ALS cases are familial with a Mendelian pattern of inheritance, and the remaining 90% of ALS cases are sporadic (2). The median survival time of ALS patients is only 3-5 years from the onset of symptoms with peak age of onset being 58-63 for sporadic ALS and 47-52 for familial ALS (1, 3, 4). The number of ALS cases across globe is expected to increase by ~70% in the next two and half decades, creating enormous potential social and health burden in the coming years (5). Both genetic and environmental factors contribute to the pathogenesis of ALS. Several disease genes have been identified to be associated with familial ALS and sporadic ALS. The familial ALS associated genes include *SOD1, TARDBP, FUS, ANG* and *OPTN* (1), each individually accounts for 20%, 5-10%, 5% and 1% of familiar ALS cases, respectively. The sporadic ALS associated genes include *ABCG1, C9orf72, CPNE4, SARM1, MOBP, ITPR2, DPP6* and *UNC13A*, with all genetic variants explaining a total of 8.5% of ALS phenotypic variation (6, 7). Besides genetic factors, metabolic traits or environmental factors such as smoking, exposure to pesticides, lead, organic toxins, or electromagnetic radiation, and socioeconomic status are also associated with ALS (8-15).

Among the various genetic and environmental factors that have been identified to be associated with ALS, the association between blood lipid metabolites and ALS has attracted considerable recent attention. The associations between lipids and ALS are strong and are comparable in strength to many previously identified key ALS risk factors (Table S1). Lipids are the major source of energy for muscles, and muscle energy alteration is frequently observed in ALS patients (8, 16-19). Specifically, ALS patients suffer from high prevalence of malnutrition, increased resting energy expenditure, and weight loss, all of which affect the quality of life for ALS patients. Previous observational studies have shown that ALS patients frequently have dyslipidemia (8). Dyslipidemia is characterized by abnormal levels of high density lipoprotein (HDL), low density lipoprotein (LDL), total cholesterol (TC) and triglycerides (TG). The positive association between dyslipidemia and ALS suggests that high non-HDL lipid levels may play a protective role in the progression of ALS. Consistent with the human observational studies, studies with ALS mice models have also shown that the overall survival of ALS mice is reduced under caloric restriction (20). However, the relationship between dyslipidemia and ALS has become controversial in recent years. Specifically, conflicting results have been reported for basal serum lipid levels, the cause of dyslipidemia, and the relationship between serum lipid levels and ALS disease progression (21). For example, many follow-up observational studies of ALS did not observe any association between dyslipidemia and ALS (22-24). In addition, some studies have shown that the patients of ALS often have hypolipidemia — which is mainly characterized by low LDL levels — in both male and female ALS patients (25, 26). The association between hypolipidemia and ALS is further confirmed in a mice ALS model (21). The conflicting results on the relationship between lipid levels and ALS may be partly due to the relatively small sample sizes employed in previous studies and partly due to uncontrolled confounding factors that are inevitable in observational studies.

Understanding the causal impact of lipids on ALS is important from both medical and public health perspectives. Because lipids are modifiable factors, a better characterization of the causal effects of lipids on ALS can facilitate the development of effective strategies for reducing ALS occurrence and the associated society burden, for treating ALS disease progression and for improving the quality of life for ALS patients. However, determining the causal impact of lipids on ALS through traditional randomized controlled trial (RCT) studies is challenging, as such studies necessarily require long-term follow-ups, are expensive, and often times, are unethical to carry out (27). Therefore, it is desirable to determine the causal relationship between lipids and ALS through observational studies. A powerful statistical tool to examine causal relationship and estimate causal effects in observational studies is Mendelian randomization. Mendelian randomization adapts the commonly used instrumental-variable analysis approach developed in the field of causal inference to settings where genetic variants [i.e. single nucleotide polymorphisms (SNPs)] are served as instruments (28, 29). In particular, Mendelian randomization employs genetic variants as proxy indicators (i.e. instrumental variables) for the modifiable exposure (i.e. lipids) and uses these genetic variants to assess the causal effect of the exposure on the outcome variable of interest (i.e. ALS) (Supplementary Material Fig. S1) (30).

Compared with alternative approaches for causal inference, Mendelian randomization has the following desirable advantages: (**i**) because genetic variants are measured with high accuracy and can potentially capture long-term exposure effects, the results of Mendelian randomization analysis are often not susceptible to estimation bias due to measurement errors that are commonly encountered in randomized clinical studies (31); (**ii**) because the two alleles of a SNP are randomly segregated during gamete formation and conception under Mendel’s law and because such segregation is independent of many known or unknown confounders, the results from Mendelian randomization are also less susceptible to reverse causation and confounding factors compared with other study designs (32). As a result, Mendelian randomization has become a popular and cost-effective analytic tool for causal inference in observational studies, alleviating the need to record and control for all possible confounding factors present in these studies (33).

Here, we perform a two-sample summary data Mendelian randomization study to comprehensively investigate the causal relationship between lipid levels and sporadic ALS. The primary goal of our study is to investigate whether HDL, LDL, TC or TG has any causal effect on ALS and whether such causal relationship holds in two different human populations that include both the European and East Asian populations. ALS incidence rates vary greatly across different populations and can be two to several times lower in the East Asian population than in the European population (4, 7, 34-36). Therefore, examining the causal relationship between lipids and ALS in both European and East Asian populations can shade light on the similarity or difference in the genetic underpinning of ALS in different populations, facilitating coordination of ALS presentation strategies across nations. To investigate the causal effects of lipids on ALS, we carry out one of the largest and most comprehensive Mendelian randomization analyses to date by using summary statistics obtained from large scale genome-wide association studies (GWASs) with ~190,000 individuals for lipids and ~42,000 individuals for ALS in the European population as well as ~130,000 individuals for lipids and ~4,100 individuals for ALS in the East Asian population.

## Results

### Causal effects of HDL, LDL, TC and TG on ALS in the European population

We first selected SNPs that can serve as valid instrumental variables for each of the four lipid traits (HDL, LDL, TC and TG) in the European population based on association summary statistics from the Global Lipids Genetics Consortium (GLGC) study (37). The GLGC study is the largest GWAS meta-analysis performed thus far on four lipid traits using 188,577 individuals of European ancestry. From the GLGC study, we obtained a total of 88, 78, 86 and 54 index SNPs to serve as instrumental variables for HDL, LDL, TC and TG (Supplementary Material, Tables S2-S5), respectively. These SNPs were obtained through the clumping procedure of PLINK (38) by following (39), are independent of each other, and are all associated with the corresponding lipid trait at the genome-wide significance level of *p* = 5.00E-8. The selected SNPs in total explained 6.15%, 8.75%, 8.05% or 4.92% of phenotypic variance (PVE) for HDL, LDL, TC or TG; respectively. Importantly, the *F* statistics for all these SNPs are above 10 (Supplementary Material, Tables S2-S5), suggesting that the selected SNPs have sufficiently strong effect sizes to serve as valid instruments and that weak instrument bias is unlikely to occur (40, 41).

With the identified instruments, we performed Mendelian randomization analyses to investigate the causal effects of each of the four lipid traits on ALS. To do so, we obtained association summary statistics from an ALS GWAS in the European population for the selected instrumental variables of the lipid traits. The ALS GWAS consists association results based on 41,689 individuals of European ancestry (7). Among the SNPs that were selected as instrumental variables, we removed SNPs that are associated with ALS with a marginal *p* value below the 0.05 level after Bonferroni correction (i.e. *p* < 0.05/*k*, where *k* is the number of the instrumental variable-savailable for the given lipid trait). With the remaining instruments, for each lipid in turn, we applied both the fixed-effects and the random-effects versions of the inverse variance weighted (IVW) methods to perform Mendelian randomization analysis. These IVW methods pool results from individual SNPs to produce an estimate of the overall causal effect of each lipid on ALS. Because there is no evidence of effect size heterogeneity across SNP instruments for any of the four lipid traits (the *p* values for these Q statistics are 0.017, 0.174, 0.161 and 0.123 for HDL, LDL, TC and TG respectively; and I^2^ statistics are 25.0%, 11.9%, 12.1% and 17.1% for HDL, LDL, TC and TQ respectively) and because the random-effects IVW yielded almost identical results to the fixed-effects IVW method (Fig. 1A), we report the results mainly from the fixed-effects IVW method in the following paragraph.

**Figure 1.**
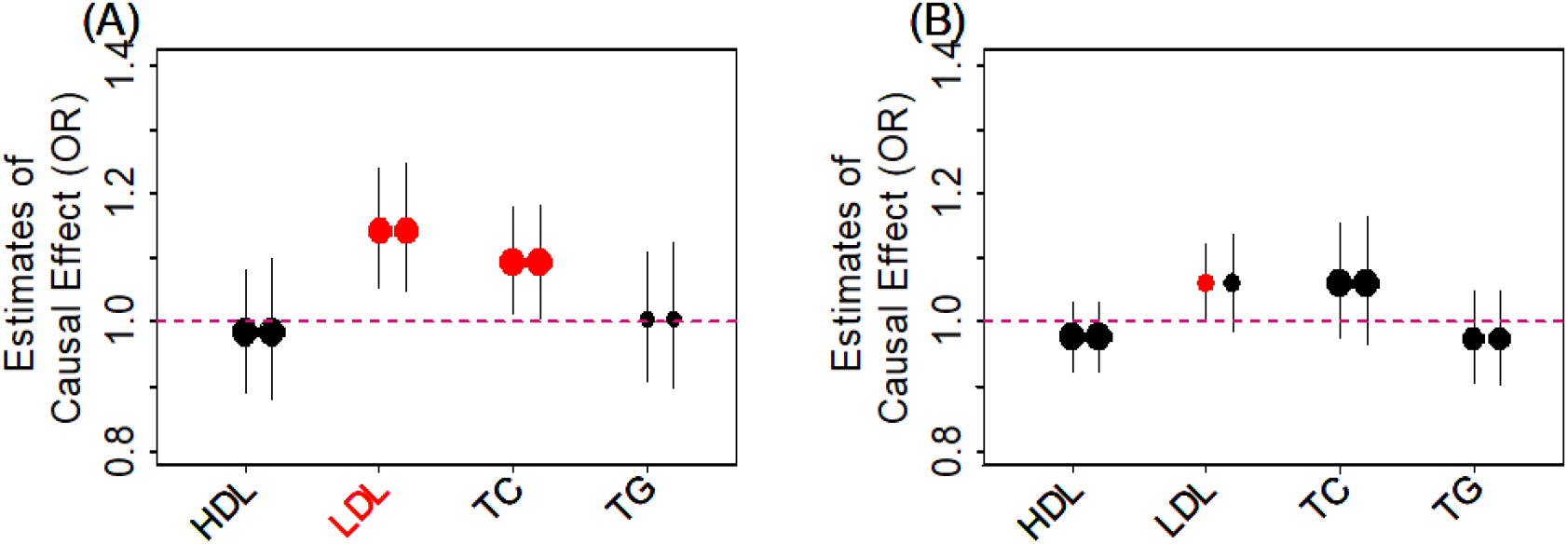
Causal effect estimates and 95% confidence intervals for four lipids on ALS. Estimates are shown in both the European population (**A**) and the East Asian population (**B**). In both plots, center of the dot represents estimated causal effects and the magnitude of the dot is in proportion to the number of the instrumental variables used in the analysis. Causal effects with a p value below a nominal level of 0.05 are highlighted in red. Name of the traits that have a causal effect p value below a Bonferroni corrected threshold (p < 0.05/4) are further colored in red on the x-axis. For each lipid, the left dot shows results using the fixed-effects IVW method and the right dot shows causal results using the random-effects IVW method.

We found that, among the four lipid traits (Fig. 1A), only LDL is causally associated with ALS at a Bonferroni corrected p-value threshold (0.05/4 = 1.25E-2). Specifically, the OR of ALS per one standard deviation (SD) increase of LDL is estimated to be 1.14 (95% CI 1.05 - 1.24, *p* = 1.38E-3), suggesting that high LDL level is causally associated with an increased risk of ALS. Among the other lipid traits, we found that TC is causally associated with ALS at a nominal p-value threshold of 0.05: the OR of ALS per one SD increase of TC is estimated to be 1.09 (95% CI 1.01 - 1.18, *p* = 2.53E-2). In contrast, the OR of ALS per one SD increase of HDL is 0.98 (95% CI 0.89 - 1.08, *p* = 0.716) and the OR of ALS per one SD increase of TG is 1.00 (95% CI 0.91 - 1.11, *p* = 0.948); both are not statistically significant. The causal effect estimates above are obtained based on the unit of mg/dL; the ORs of ALS for HDL, LDL, TC and TG are respectively estimated to be 0.51 (95% CI 0.01 - 18.63, *p* = 0.716), 135.71 (95% CI 6.70 - 2747.14, *p* = 1.38E-3), 24.67 (95% CI 1.49 - 408.87, *p* = 2.53E-2) and 1.13 (95% CI 0.03 - 47.98, *p* = 0. 948) if the unit of mmol/L were employed. Note that, the p values are unchanged and the difference between the two types of estimates are due to the scale difference of the two types of units (i.e. mg/dL vs. mmol/L).

Finally, to examine result robustness, instead of using the SNP instruments identified by the clumping procedure, we performed an additional Mendelian randomization analysis using independent association SNPs reported in the original study (37) as instruments (71 instruments for HDL, 58 for LDL, 74 for TC, and 40 for TG). With the new set of instruments, we obtained almost identical estimation results. For example, with fixed-effects IVW, the ORs of ALS per one standard deviation (SD) for HDL, LDL, TC and TG are estimated to be 0.97 (95% CI 0.87 - 1.07, *p* = 0.525), 1.11 (95% CI 1.01 - 1.24, *p* = 4.51E-2), 1.09 (95% CI 0.99 - 1.19, *p* = 7.76E-2) and 1.0 (95% CI 0.90 - 1.11, *p* = 0.990).

Because we only detected a non-zero causal effect of LDL on ALS, we examined whether the lack of detectable non-zero causal effects of the other lipid traits are due to a lack of statistical power for these traits in the Mendelian randomization analysis. To do so, for each lipid in turn, we performed simulations and power analyses using 41,689 individuals. We estimated the statistical power to detect an OR ratio of 1.10 (or 0.90) in the risk of ALS per unit change of the lipid following the method in (42). The statistical power based on a Bonferroni adjusted threshold to detect such amount of causal effects from HDL, LDL, TC and TG on ALS are estimated to be 44%, 62%, 57% and 34%, respectively. The power results suggest that we would have comparable and reasonable power across the four lipids.

### Causal effects of HDL, LDL, TC and TG on ALS in the East Asian population

We next estimated the causal effects of HDL, LDL, TC and TG on ALS in the East Asian population. To do so, we obtained association summary statistics from an East Asian lipids GWAS with 162,255 Japanese individuals (43) and corresponding association summary statistics from another East Asian ALS GWAS on 4,084 Chinese individuals (4). We first obtained a set of 31, 22, 32 and 26 lipid associated SNPs to serve as instrumental variables for HDL, LDL, TC and TG; respectively (Supplementary Material, Tables S6-S9). The estimated PVE by the selected SNPs are 7.75%, 3.89%, 2.67% and 5.02% for HDL, LDL, TC and TG; respectively. All selected SNPs have an *F* statistic above 10 and are thus considered as strong instruments (Supplementary Material, Tables S6-S9) (40, 41).

With the identified instruments, we performed Mendelian randomization analyses using both the fixed-effects and random-effects IVW methods to investigate the causal effects of each of the four lipid traits on ALS. Like what we observed in the European data, we did not detect any SNP effect size heterogeneity on any of the four lipids in the Asian population: the *p* values for the Q statistics are 0.494, 0.066, 0.161 and 0.769 for HDL, LDL, TC and TG; respectively; the I^2^ statistics are 0%, 27.3%, 14.9% and 0% for HDL, LDL, TC and TG; respectively. In addition, the results from both the fixed-effects and random-effects IVW analysis are largely consistent with each other (Fig. 1B). Specifically, in the fixed-effects IVW analyses, we found that high LDL is causally associated with an increased risk of ALS at the significance level of 0.05 (Fig. 1B). The OR of ALS per one SD increase of LDL is estimated to be 1.06 (95% CI 1.0 - 1.12, *p* = 0.044). In contrast, the ORs of ALS per one SD increase of HDL, TC and TG are 0.97 (95% CI 0.92 - 1.03, *p* = 0.361), 1.06 (95% CI 0.98 - 1.15, *p* = 0.165), and 0.97 (95% CI 0.90 - 1.05, *p* = 0.470), respectively.

Next, we examined whether the lack of detectable non-zero causal effects of the other non-LDL lipid traits in the East Asian population were due to a lack of statistical power for these traits. To do so, for each lipid in turn, we performed simulations and power analyses using 4,084 individuals. We estimated the statistical power to detect an OR ratio of 1.10 (or 0.90) in the risk of ALS per unit change of the lipid. The statistical powers based on a Bonferroni adjusted threshold to detect such amount of causal effects from HDL, LDL, TC and TG on ALS were estimated to be 4%, 3%, 2% and 3%, respectively (or 13%, 8%, 8% and 10% based on a nominal p value threshold of 0.05). Simulation results suggest that we have a small amount of power to detect non-zero lipid effects due to the small sample size in the Asian ALS GWAS. Therefore, larger sample size is likely needed to validate our causal effect estimates in the East Asian population.

Finally, we note that we did not observe statistically significant causal effect size heterogeneity between the two populations for any of the lipid on ALS. Specifically, the causal effect estimates of each lipid on ALS from the two populations share the same directionality and their 95% confidence intervals are overlapped (*p* = 0.115 with a *z*-test). Therefore, we used the inverse-variance weighted meta-analysis method to combine the causal effect estimates from both European and East Asian populations to obtain a joint causal effect estimate. By assuming identical causal effects of lipids on ALS in both populations, we estimated the ORs of ALS per one SD increase of HDL, LDL, TC and TG to be 0.98 (95% CI 0.93 - 1.02, *p* = 0.165), 1.08 (95% CI 0.1.03 - 1.14, *p* = 2.54E-4), 1.08 (95% CI 1.02 - 1.14, *p* = 4.81E-3) and 0.98 (95% CI 0.93 - 1.04, *p* = 0.292), respectively. The combined results confirm our main conclusion that LDL is positively causally associated with ALS.

### Sensitivity analyses confirm the causal effects of LDL on ALS

We now validate the causal relationship between LDL and ALS inferred in both European and East Asian populations through various sensitivity analyses. First, to guard against the possibility that the associated instruments for one lipid may exhibit their causal effects on ALS through the other lipids, we performed Mendelian randomization for each lipid using lipid-specific instruments (Supplementary Material, Fig. S2). These lipid-specific instruments were obtained by removing those instruments that were also selected as instruments for the other three lipids (i.e. based on the same clumping strategy with a significance level of *p* = 5.00E-8). The Mendelian randomization results using these lipid-specific instruments are consistent with the main results in both European and East Asian populations (Supplementary Material, Table S10).

Next, for LDL, we also excluded potential pleiotropic instrumental effects by performing a residual-based (or joint) multivariable Mendelian randomization analysis with HDL and TG treated as covariates (44, 45). Note that we cannot include TC in addition to HDL and TG as covariate in the model because LDL level can be completely determined by the other three lipids (46) and is highly correlated with TC (correlation coefficient *r* = 0.992 and 0.787 in the European and East Asian populations). The multivariate Mendelian randomization analysis tests for the causal effects of LDL on ALS while controlling for the other lipid effects, thus providing a way to control for instrumental pleiotropic effects on other lipids. The estimation results from the residual-based (or joint) multivariable Mendelian randomization analysis are again consistent with those in the main text (Supplementary Material, Table S10), supporting the observation that LDL has a positive causal effect on ALS. Specifically, after removing the effects of HDL and TG and applying the residual-based multivariable regression, the OR of ALS per one SD increase of LDL is estimated to be 1.13 (95% CI 1.03 - 1.23, *p* = 8.84E-3) in the European population, albeit less significant in the East Asian population presumably due to the smaller number of instruments used there (Supplementary Material, Table S10). The results above imply that the causal effect of LDL on ALS is unlikely acted through the other lipids.

Next, we examined whether there exist potential SNP instrument outliers and whether these outliers influence Mendelian randomization analysis results. To do so, we visualized all SNP instruments by plotting the SNP effect sizes on LDL versus their effect sizes on ALS in Fig. 2. In the European population, three SNPs (rs11591147, rs7254892 and rs12721109) have effect sizes on LDL larger than 0.40 and these SNPs appear to be potential outliers (Fig. 2A). However, these potential outliers do not influence the causal effect estimation in the Mendelian randomization analysis. Indeed, after removing these three SNPs, the OR of ALS per one SD increase of LDL is estimated to be 1.15 (95% CI 1.05 - 1.25, *p* = 1.04E-3) (Fig. 2B), almost identical to that obtained using all instruments (i.e. OR = 1.14, 95% CI 1.05 - 1.24, *p* = 1.38E-3). In the East Asian population, one SNP, rs769446, has a large effect size of −0.51 (95% CI −0.54 - −0.49, *p* = 2.98E-322) on LDL and appears to be an outlier on the effect size scatter plot (Fig. 2C). After removing this SNP, the OR of ALS per one SD increase of LDL is estimated to be 1.24 (95% CI 1.11 - 1.39, *p* = 1.54E-4) (Fig. 2D), which is larger and more significantly different from zero than the estimate obtained using all instruments (i.e. OR = 1.06, 95% CI 1.00 - 1.12, *p* = 0.044). Therefore, even though rs769446 may be a potential outlier, it only supports our main conclusion that LDL has a statistically significantly positive causal effect on ALS in the East Asian population.

**Figure 2.**
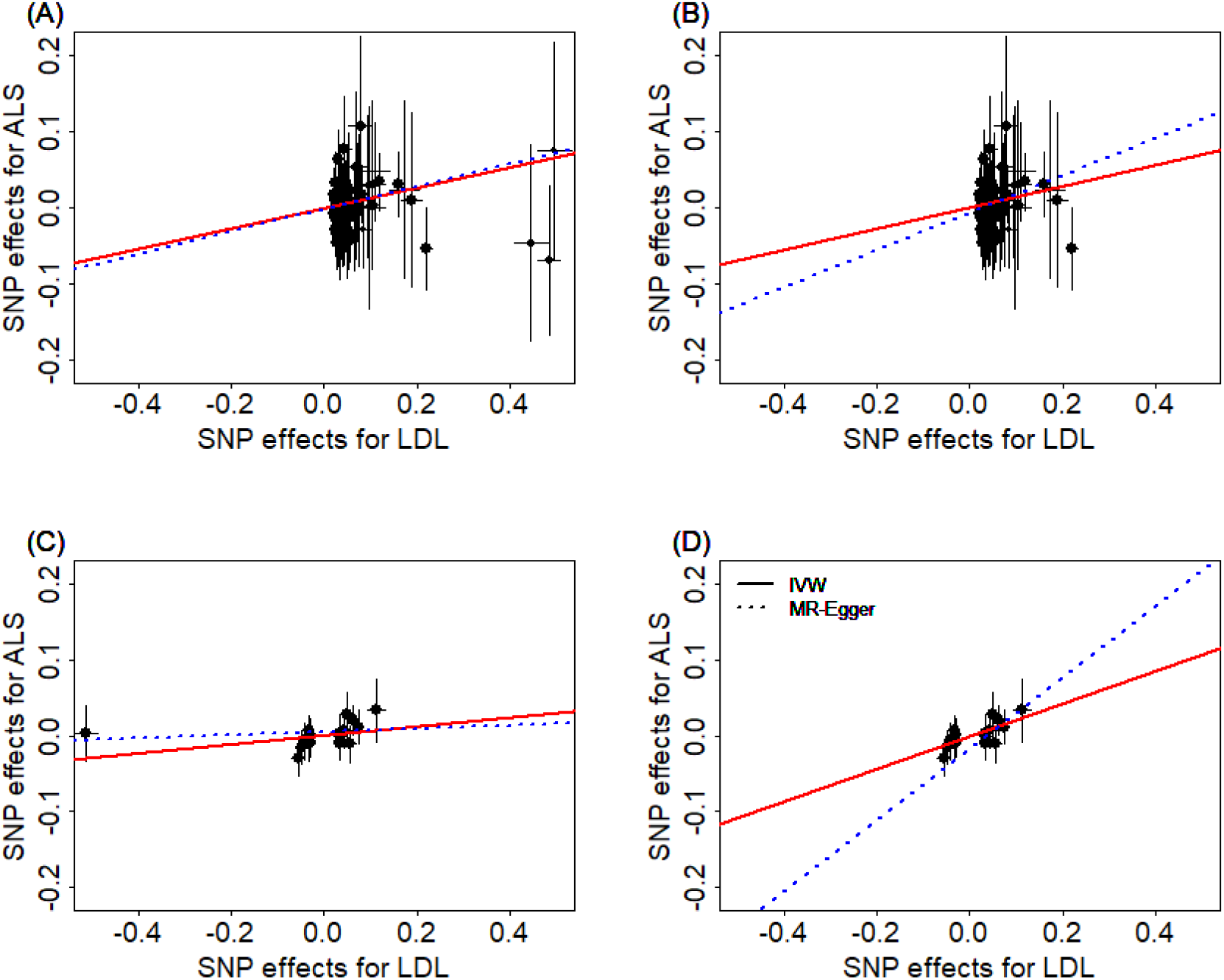
Relationship between the SNP effect size estimates on LDL (x-axis) and the corresponding effect size estimates on ALS (y-axis) for SNPs that serve as instrumental variables. Results are shown in the European population for all SNP instruments (**A**) and for the remaining instrumental SNPs after removing three potential outliers (**B**). Results are also shown in the East Asian population for all SNP instruments (**C**) and for the remaining instrumental SNPs after removing one potential outlier (**D**). On each plot, 95% confidence intervals for the estimated SNP effect sizes on ALS are shown as vertical black lines, while the 95% confidence intervals for the estimated SNP effect sizes on LDL are shown as horizontal black lines. The vertical and horizontal red dotted lines represent zero effects. The slope of fitted lines represents the estimated the casual effects of LDL on ALS obtained using either the IVW methods (red solid lines) or the MR-Egger regression (blue dotted lines). Due to the inclusion of an intercept in the MR-Egger regression, the fitted lines by MR-Egger regression (blue dotted lines) do not necessarily pass the origin.

To formally rule out the possibility that a single instrument may strongly influence the point estimates and the statistical test results, we performed a leave-one-out (LOO) Mendelian randomization analysis. In the European population, the LOO analysis results show that no single instrument acts as an outlier and influences the causal effect estimates (Supplementary Material, Fig. S3A). In the East Asian population, the only detected outlier is rs769446 (Supplementary Material, Fig. S3B). However, as described in the previous paragraph, removing rs769446 does not influence the conclusion that LDL has a positive causal effect on ALS.

To guard against the possibility that some instruments are invalid, we conducted a Mendelian randomization analysis using the weighted median method (47). The weighted median method yields qualitatively similar results as our main analysis. In particular, the ORs of ALS per one SD of LDL are estimated to be 1.17 (95% CI 1.03 - 1.33, *p* = 1.60E-2) in the European population and 1.29 (95% CI 1.11 - 1.49, *p* = 6.29E-4) in the East Asian population (after removing the outlier rs769446), respectively. The weighted median analysis results suggest that invalid instruments unlikely bias our main results.

To guard against the possibility of horizontal pleiotropy of these instruments, we performed Mendelian randomization using the MR-Egger regression (48, 49). The results from the MR-Egger regression analysis are again largely consistent with our main results. For example, in the European population, MR-Egger regression estimates the OR of ALS per one SD increase of LDL to be 1.16 (95% CI 1.02 - 1.32, *p* = 0.026) when all instruments are used. When the three SNP outliers (rs11591147, rs7254892 and rs12721109) are removed, MR-Egger regression estimates the OR of ALS per one increase SD of LDL to be 1.23 (95% CI 1.05 - 1.45, *p* = 9.19E-3). In the East Asian population, MR-Egger regression estimates the OR of ALS per one SD increase of LDL to be 1.02 (95% CI 0.94 - 1.11, *p* = 0.645) when all instruments are used. When the SNP outlier (rs12927205) is removed, MR-Egger regression estimates the OR of ALS per one SD increase of LDL to be 1.60 (95% CI 1.25 −2.06, *p* = 2.41E-4). Examining the intercept estimates from MR-Egger regression also leads to the same conclusion that horizontal pleiotropy unlikely influence results in either population. For example, in the European population, the MR-Egger regression intercept was estimated to be −0.001 (95% CI −0.009 - 0.007, *p* = 0.767) when all instruments were used, and was estimated to be −0.005 (95% CI −0.015 - 0.004, *p* = 0.289) when three instrument outliers were removed. In the East Asian population, the MR-Egger regression intercept was estimated to be 0.006 (95% CI −0.003 - 0.014, *p* = 0. 180) when all instruments were used, and was estimated to be −0.016 (95% CI −0.029 - −0.003, *p* = 0.020) when one instrument outlier is removed. Furthermore, funnel plots also display symmetric pattern of effect size variation around the causal effect point estimate (Fig. 3). Together, funnel plots and MR-Egger regression analyses suggest that horizontal pleiotropy unlikely bias our results.

**Figure 3.**
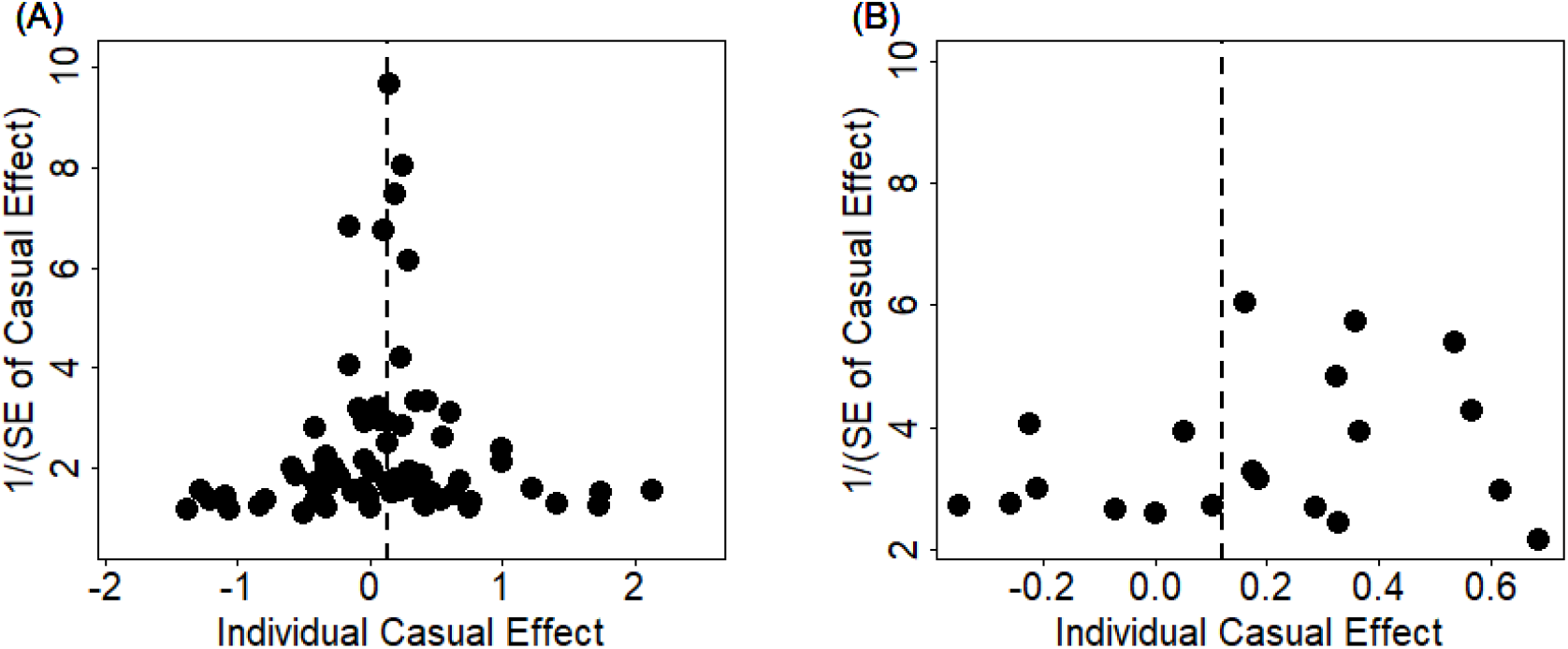
Funnel plot displays causal effect estimates of LDL on ALS obtained using individual SNP instruments in the European population (**A**) and the East Asian population (**B**). Center of the dots represents the estimated causal effect (log OR) for each instrumental variable. The vertical dotted lines represent the estimated causal effect obtained using all instrumental variables.

To examine the possibility that our results are driven by reverse causality, we performed IVW analysis in the reverse direction to examine the causal effects of ALS on LDL. In total, we identified 5 and 1 SNPs to serve as instruments for ALS in the European and East Asian populations, respectively. Results show that there is no detectable causal effect of ALS on LDL: the causal effect of ALS on LDL is estimated to be 0.003 (95% CI −0.022 - 0.027, *p* = 0.834) in the European population and 0.094 (95% CI −0.145 - 0.334, *p* = 0.439) in the East Asian population, respectively.

Finally, we note that several previous studies have shown that the ratio of LDL/HDL, instead of LDL alone, is also correlated with ALS (8, 9). In our results, we have shown that HDL is negatively associated with ALS (though not statistically significant) while LDL is positively associated with ALS, suggesting that the ratio of LDL/HDL is likely positively associated with ALS. While it is tempting to directly examine the causal effect of the ratio of LDL/HDL on ALS, we acknowledge that it would be challenging to do so because there is currently no GWAS summary statistics available for the ratio of LDL/HDL. Instead of directly investigating the causal effects of LDL/HDL on ALS, we indirectly examined this relationship by estimating the causal effect of LDL on ALS while controlling for HDL. Intuitively, if LDL/HDL is high for an individual, then the residual value of LDL after controlling for HDL would also be high for the individual. Therefore, if LDL/HDL is associated with ALS status, then LDL after controlling for HDL will also likely be associated with ALS. With multivariable Mendelian randomization analysis (50), we found that, after removing the effect of LDL/HDL, the OR of ALS per one SD increase of LDL is nearly unchanged and is estimated to be 1.13 (95% CI 1.04 - 1.23, *p* = 7.58E-3) in the European population and increases to 1.26 (95% CI 1.13 - 1.41, *p* = 5.65E-4) in the East Asian population.

## Discussion

We have presented a comprehensive Mendelian randomization analysis to investigate the causal relationship between lipids and ALS using summary statistics from large GWASs. The large sample size used in our study allows us to fully establish a positive causal effect of the modifiable factor LDL on ALS in both European and East Asian populations. The inferred causal relationship between LDL and ALS is robust with respect to the choice of statistical methods and is carefully validated through various sensitivity analyses. Overall, our study provides important evidence supporting the causal role of higher LDL on increasing the risk of ALS.

The positive causal effect of LDL on ALS suggests that future development of strategies to reduce LDL levels would likely reduce disease burden of ALS (51). LDL is a modifiable risk factor whose levels can be reduced through various intervention strategies. For example, dietary changes such as increased fiber consumption, increased phytosterol consumption, and increased nut consumption can all lead to a reduction in LDL levels (52). Restrictions in dietary cholesterol intake, restrictions in high-carbohydrate diets, and restrictions in trans-fatty acid consumption can also reduce LDL levels (53). Beside lifestyle and dietary changes, lowering LDL can be achieved by drug-based therapy. For example, various drugs such as cholestyramine, niacin, ezetimibe, statins can each significantly reduce LDL levels (54). Future development of LDL reduction strategies and the making of public policies to promote such strategies would likely lead to a reduction of ALS disease burden in the society.

Our results dovetail a recently published Mendelian randomization study which relies on instrumental variables associated with both LDL and TC to investigate their joint causal effect on ALS (55). The estimated causal effect of LDL on ALS in our study is consistent with the estimated causal effect of LDL and TC on ALS in Chen et al. (55) both in terms of magnitude and directionality (OR = 1.14 for LDL on ALS in our study vs OR=1.16 for LDL in Chen et al; OR = 1.09 for TC on ALS in our study vs OR=1.15 for TC on ALS in Chen et al. for the European population). However, compared with (55), our study has several unique advantages. First, the samples used in our study to select the instruments for lipids are proximately two times larger than that in Chen, et al. (55) (i.e. ~190,000 vs. ~100,000). Therefore, we are able to identify more and stronger instruments than that in Chen, et al., leading to a potential power gain of Mendelian randomization analysis (30, 41, 56, 57). Second, unlike Chen et al., we have performed extensive sensitivity analyses (e.g. leave-one-out, median-based and multivariable analyses, Egger regression, and guarding against horizontal pleiotropic effects etc. (58)) to ensure the robustness of our results and to guard against any possible model assumption violation in the Mendelian randomization analysis. Third, by examining both European and East Asian populations, we are able to establish a causal effect of higher LDL on increasing risk of ALS in both populations. The inferred causal relationship between LDL and ALS in multiple populations suggests that strategies to reduce LDL levels have the potential to lower ALS in multiple populations and ethnic groups.

The causal relationship between lipids and ALS identified in the European and East Asian populations are inferred using a different set of instruments in the two populations. Indeed, the overlap between the index SNPs used in the European population and the index SNPs used in the East Asian population is small: only 2, 1, 2 and 1 index SNPs are overlapped between the two populations for HDL, LDL, TC and TG; respectively. However, most index SNPs from one population are in linkage disequilibrium with index SNPs from another population, suggesting similar genetic architectures underlying lipid traits in different populations. In addition, we found that the causal effects of LDL and TC on ALS estimated in the European population are comparable with those estimated in the East Asian population (e.g. 1.14 vs. 1.06 for LDL, and 1.09 vs. 1.06 for TC), allowing us to combine causal effect estimates from the two populations via meta-analysis. The similarity in the causal effects of LDL on ALS in the two populations suggests potential commonality of ALS etiology in multiple populations. However, we also recognize that there remains substantial unexplained diversity underlying ALS etiology in the two populations in terms of, for example, the different prevalence rates of ALS. Future studies are needed to understand the genetic and environmental basis underlying ALS prevalence difference across populations.

Our results on the causal effects of lipids on ALS in the European population are obtained using a European lipids GWAS and a European ALS GWAS. However, our results in the East Asian population are obtained using a Japanese lipids GWAS and a Chinese ALS GWAS. Because our East Asian results are based on using two different ethnic groups, one might concern that the genetic difference between the two ethnic groups may bias our causal effect estimates. However, it is well known that the major East Asian ethnic groups (e.g. Japanese, Chinese and Korean) are genetically closely related to each other with similar genetic architectures (59, 60). Indeed, recent studies have shown that the genetic difference among multiple East Asian ethnic groups is comparable to the genetic difference among multiple European ethnic groups, both in terms of time to the most recent common ancestor (TMRCA) or divergence time (61). Consequently, GWAS meta-analyses in East Asian are commonly performed by combining association results from multiple ethnic groups (62-66), as we also do here. Therefore, our causal effect estimation results in the East Asian population are unlikely biased by the genetic difference between the two ethnic groups. However, to further alleviate concerns, we have conducted additional Mendelian randomization analyses to estimate the causal effects of lipids on ALS using a Chinese lipids GWAS (Supplementary Material, Tables S11-12) (67), which has a smaller sample size than the Japanese lipids GWAS (23,083 vs. 94,356). The results are largely consistent with the results we have presented in the main text (Supplementary Material, Fig. S4). For example, the OR of ALS per one SD increase of LDL is estimated to be 1.18 (95% CI 1. 04 - 1.35; *p* = 9.44E-3) using the Chinese lipids GWAS, close to what we have obtained using the Japanese lipids GWAS. The results based on the Chinese lipids GWAS further confirm a positive causal effect of LDL on the risk of ALS in the East Asian population.

Like many other Mendelian randomization analyses, our results are not without limitations. First, we emphasize again that the validity of Mendelian randomization analyses relies on three key modeling assumptions listed in Supplementary Material Fig. S1 (30, 33, 57). While we have been extremely careful in selecting instrumental variables to satisfy these assumptions and have performed various sensitivity analyses to guard against various model assumption violations, we still would like to follow the standard practice of Mendelian randomization analysis to caution against the over-interpretation of the causality results obtained in the present study (29, 41, 56, 68, 69). Second, like other Mendelian randomization applications, we have assumed a linear relationship between lipids and ALS in the Mendelian randomization model. While a linear relationship can be viewed as a first order approximation to any non-linear relationship, we acknowledge that modeling a linear relationship can be suboptimal in terms of power if the true relationship is non-linear. Third, due to the use of GWAS summary statistics, we cannot carry out stratified analyses by gender to estimate the causal effects of lipids on ALS in males and females separately. Therefore, we are unable to estimate or validate previously estimated gender-specific effects of lipids on ALS in observational studies (25, 26).

## Materials and Methods

We provide an overview of the materials and methods used in the present study here, with details available in Supplementary Material.

### GWAS data sources

In the European population, we obtained association summary statistics for SNPs in terms of their marginal effect size estimates and standard errors for each of the four lipid traits (HDL, LDL, TC and TG) from the Global Lipids Genetics Consortium (GLGC) study (Supplementary Material, Table S13) (37). The GLGC study is the largest GWAS performed to date on lipid traits and is a meta-analysis that analyzed ~2.5 million genotyped and imputed SNPs on 188,577 individuals. To examine the causal effect of lipids on ALS in Europe, we also obtained association summary statistics from an European ALS GWAS that analyzed ~8.7 million genotyped and imputed SNPs on 41,689 individuals (14,791 cases and 26,898 controls) (Supplementary Material, Table S14) (7). In the East Asian population, we obtained association summary statistics for lipid traits from a Japanese lipids GWAS. The Japanese lipids GWAS analyzed a total of ~6.0 million genotyped and imputed SNPs, on 70,657 individuals for HDL, 72,866 individuals for LDL, 128,305 individuals for TC and 105,597 individuals for TG (Supplementary Material, Table S15) (43). To examine the causal effect of lipids on ALS in East Asia, we also obtained association summary statistics from a Chinese ALS GWAS that analyzed ~6.6 million genotyped and imputed SNPs on up to 4,084 individuals (1,234 cases and 2,850 controls) (4).

### Instrumental variable selection

In the European population, from the GLGC study (37), we selected 88, 78, 86 and 54 independent index association SNPs to serve as instrumental variables for HDL, LDL, TC and TG (Supplementary Material, Tables S2-S5), respectively. These index SNPs were obtained using the clumping procedure of the plink software (38) following previous work (39). To estimate the causal effect of each of the four lipids on ALS, we also extracted association summary statistics with ALS for these index SNPs from the European ALS GWAS (7). In the East Asian population, from the Japanese lipids GWAS (43), we selected 30, 21, 31 and 26 index association SNPs to serve as instrumental variables for HDL, LDL, TC and TG (Supplementary Material, Tables S6-S9), respectively. To estimate the causal effect of each of the four lipids on ALS, we also extracted association summary statistics of these index SNPs for ALS from the Chinese ALS GWAS (4).

### Estimation of causal effect with Mendelian randomization

In either population, we excluded index SNPs that show pleiotropic associations for ALS to ensure the validity of Mendelian randomization analysis (30, 56). However, none of the index SNPs show pleiotropic associations with ALS in both populations in our main text analyses, thus we kept all of them in the analysis. For each index SNP in turn, we estimated the proportion of phenotypic variance for the corresponding lipid trait explained by the index SNP using summary statistics following the method presented in (70). We further computed *F* statistic to ensure strong instruments (40, 41). We then performed Mendelian randomization analysis for one lipid at a time to estimate its causal effect on ALS in terms of the odds ratio (OR) for one-unit change of lipid level, where the unit is defined as standard deviation (SD) of the trait, estimated to be 15.1 mg/dL for HDL, 39.0 mg/dL for LDL, 40.6 mg/dL for TC and 79.3 mg/dL for TG. Our two-sample Mendelian randomization analyses (48, 71) were performed using either the fixed-effects version and the random-effects version of the inverse-variance weighted (IVW) methods (41, 72). Compared with the fixed-effects version, the random-effects version of IVW accounts for causal effect size heterogeneity across instruments and often yield more conservative results. Besides applying the random-effects version of IVW, we also computed both Q and I^2^ statistics to directly measure effect size heterogeneity in the data (71, 73).

### Sensitivity and other complementary analyses

In both populations, to ensure results robustness and guard against various modeling misspecifications, we performed extensive sensitivity analyses. These analyses include: (**i**) Mendelian randomization analysis using lipid-specific instruments to guard against the possibility that some associated instruments for one lipid may exhibit their causal effects on ALS through the other lipids; (**ii**) leave-one-out (LOO) cross-validation analysis to examine whether there are instrumental outliers and whether these outliers affect results (39); (**iii**) median-based method to directly exclude outlier influence (47); (**iv**) MR-Egger regression to examine the assumption of directional pleiotropic effects (48, 49); (**v**) reverse causation analysis to exclude the possibility that ALS affects lipids; (**vi**) in addition to the sensitivity analyses, we also performed a complementary Mendelian randomization analysis to indirectly test the causal effect of the ratio of LDL/HDL on ALS (8, 9) by examining the causal effect of LDL on ALS while controlling for LDL/HDL (44, 50). We also considered to remove the potential mediation effect of HDL and/or TG by performing residual-based multivariable Mendelian randomization analysis as done in (44, 45).

## Supplementary Material

The details of materials and methods and other results are available in Supplementary Material.

## Abbreviations

HDL: high density lipoprotein
LDL: low density lipoprotein
TC: total cholesterol
TG: triglycerides
OR: odds ratio
ALS: Amyotrophic lateral sclerosis
RCT: randomized controlled trial
SNP: single nucleotide polymorphism
GLGC: Global Lipids Genetics Consortium
IVW: inverse-variance weighted
LOO: leave-one-out
GWAS: genome-wide association study

## Acknowledgements

We thank all the GWAS consortium studies for making the summary data publicly available and are grateful of all the investigators and participants contributed to those studies.

